# First Usutu virus detections in wild birds in Scotland, 2025

**DOI:** 10.64898/2026.06.11.731606

**Authors:** Andra-Maria Ionescu, Ben Jones, Robert C. Bruce, Fiona Mccraken, Nicholas Johnson, Charlotte Clough, Caroline Robinson, Heather Stevenson, Fiona Howie, Georgia Kirby, Meshach Lee, Kieran Killen, Davide Dominoni, Paul Baker, Emma Davies, Ryan Carmichael, Jean-Philippe Parvy, Emilie Pondeville, Heather M. Ferguson, Arran J. Folly

## Abstract

In summer 2025, several Eurasian Blackbird (*Turdus merula*) deaths were reported on the Isle of Arran in Scotland. Initial investigation included post-mortem examination, where no diagnosis was achieved. Following *Orthoflavivirus* and avian paramyxovirus testing, Usutu virus RNA was detected in two Blackbirds by reverse transcription-PCR. Phylogenetic analysis identified Usutu virus Africa 3.2 lineage which clustered closely with existing UK detections, indicating geographic expansion rather than a new incursion. Subsequent surveillance confirmed the presence of several potential mosquito vector species.

## Introduction

Usutu virus (USUV: *Orthoflavivirus*), a mosquito-borne zoonotic virus known to cause significant Blackbird population declines (1, 2), has circulated in southern mainland Europe for three decades. Recently, the virus has expanded northward into the Netherlands (3, 4), south-east England (5) and Denmark (6). This is likely facilitated by climate change, with warmer, prolonged summers increasing mosquito abundance and milder winters aiding vector survival (7). USUV has circulated in south-east England since 2020; with genomic sequencing providing evidence of establishment and subsequent autochthonous transmission (8). Retrospective surveillance detected West Nile Virus (WNV) RNA-an *Orthoflavivirus* closely related to USUV-in two mosquito pools from Nottingham, England in 2023 (9), indicating orthoflaviviruses can circulate at higher latitudes. However, prior to 2025, surveillance in birds in the United Kingdom had not detected USUV north of Cambridgeshire. Here, we report the first USUV detections in Blackbirds in Scotland.

### Wild bird *Orthoflavivirus* surveillance in the United Kingdom

Since 2002, the Animal and Plant Health Agency in collaboration with the Institute of Zoology (https://www.gardenwildlifehealth.org) and SRUC Veterinary Services has carried out vector-borne disease (VBD) surveillance in the United Kingdom. Wild bird deaths and morbidity are recorded, and carcasses submitted for post-mortem examination (PME). During the mosquito active season (April-November), targeted surveillance for orthoflaviviruses is conducted on samples collected from wild birds. In 2025, samples from 102 wild birds from Scotland were submitted for molecular testing and fifty-one samples from avian species known to be susceptible to USUV were tested.

### Isle of Arran outbreak identification and confirmation

Sporadic, isolated observations of Blackbirds showing ataxia were observed by locals and veterinarians in 2023 (n=1) and 2024 (n=2), followed by a cluster (20-22) between 17^th^ June and 28^th^ July 2025. This cluster included birds showing neurological signs or found dead in several areas of the island (SI File 1). Through surveillance funded by the Scottish Government’s Wild Bird Disease Surveillance programme, three Blackbird carcasses underwent PME and tissue harvesting at SRUC Veterinary Services; including a juvenile and adult female of normal weight and appearance and an ataxic bird that was too autolysed at necropsy to determine age or sex.

Tissue samples were sent to APHA for avian influenza and *Orthoflavivirus* testing. No avian influenza RNA was detected. Viral RNA was extracted from pooled brain and kidney samples for WNV and USUV testing. Based on RT-PCR assays, all three samples were negative for WNV RNA (10), but the juvenile and the severely autolysed Blackbirds were positive for USUV (11) (cycle threshold [*Ct*]: 34.13 and 35.15, respectively). The positive samples were confirmed using a pan-flavivirus RT-PCR (12) (*Ct* values: 32.33 and 33.25, respectively). Histopathology was not possible due to severe autolysis. Attempts to isolate the virus in Vero cells from sample homogenates were unsuccessful. Due to the clinical signs described, particularly torticollis, samples were tested for avian paramyxovirus which wasn’t detected.

### Next generation sequencing analysis

Metagenomics using Oxford Nanopore Technology was undertaken on the RT-PCR USUV positive Blackbird samples. Sequencing produced ca 330,000 reads for the juvenile and 30,000 reads for the autolysed adult bird. Through *de novo* assembly (SI File 2), two contigs for the juvenile (1521bp and 605bp) and one contig for the severely autolysed Blackbird (603bp) matched a representative USUV genome (GenBank accession number: NC_006551-Austria/Blackbird/2001). The 1521bp contig was included in phylogenetic analysis and aligned (MEGA, v.11) with the corresponding region from 59 UK and European USUV sequences obtained from GeneBank (SI File 3), and 12 additional sequences from UK detections. Sequences used were of African and European lineages. Sequence clustered closely with other UK and Danish detections of lineage Africa 3.2 (6) (Figure 1).

**Figure 1.**
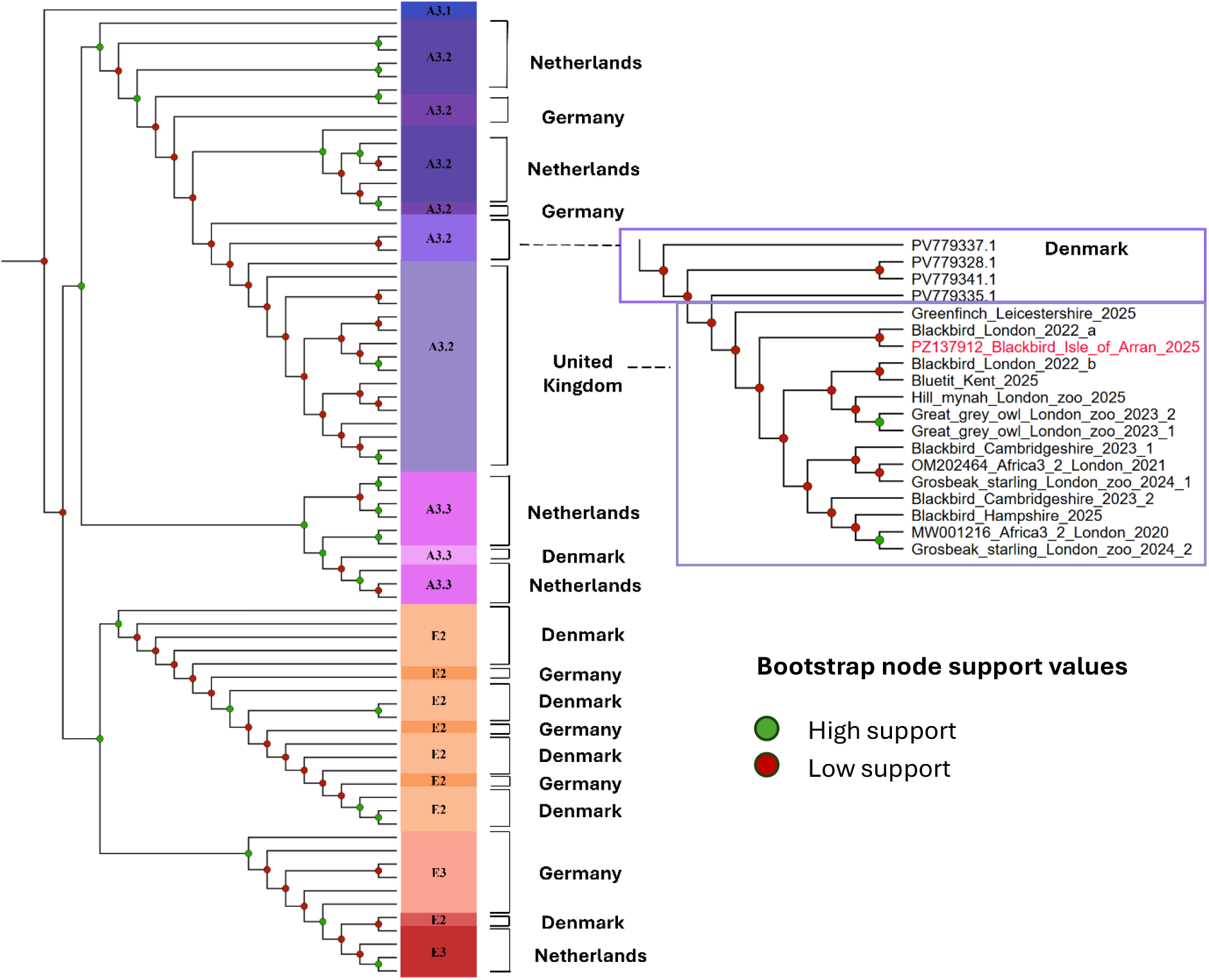
Maximum likelihood phylogenetic tree of a 1521bp USUV consensus contig from juvenile Blackbird, Isle of Arran, Scotland, July 2025, aligned against 59 genomes from GenBank of Africa 3.1-3.3 and Europe 2&3 lineages. Sequences from 12 other UK USUV detections (2022-2025) are also included, with UK clade expanded on the right-hand side of the figure. Alignment was constructed using IQ-TREE (v2.3.6). The best-fit substitution model (TNe +G4) was selected with ModelFinder. Node support values represent ultrafast bootstrap (UFBoot) percentages based on 2,000 replicates with BNNI optimisation enabled to reduce potential bootstrap overestimation. Nodes with support values greater than 75% (high support) are shown as green circles, whereas nodes with support values below 75% are shown as red circles (low support). The tree is rooted on Africa 3.1 sequence (MN122238). The 1521bp Isle of Arran sequence is highlighted in red (GenBank-PZ137912). All sequences used in the phylogenetic analysis can be found in SI File 3.

In response to observed bird deaths and clinical signs consistent with USUV, reactive surveillance of mosquitoes and Blackbirds was conducted on Arran between October 2025-Feb 2026 to identify potential vector species and assess evidence of wider avian exposure (SI File 4). All mosquito and bird samples were screened for USUV by RT-qPCR, with bird serum samples also tested for previous exposure by serology (SI File 5). Extensive searching of potential overwintering hibernacula and mosquito trapping at multiple sites yielded eight *Culex pipiens* sl (SI File 6), all confirmed as the pipiens biotype. Eighteen *Culiseta annulata*, one *Anopheles claviger* and one *Aedes detritus* were also collected (SI File 6). Blood samples from 67 birds were obtained through mistnetting: including at sites where Blackbirds with neurological signs were previously observed (SI File 6). The majority were Blackbirds (n=43), with limited numbers of 8 other species (SI File 6). All mosquito and bird samples were negative for USUV RNA, and all bird serum samples were negative for anti-flavivirus antibodies.

## Discussion

Here we report the first detection of USUV RNA in wild birds in Scotland. Phylogenetic analysis indicated that the sequence falls within USUV African lineage 3.2 and clusters with existing UK USUV detections, suggesting limited divergence from the initial introduction, consistent with expansion rather than a new introduction. Given Blackbird mortality was observed over five weeks, findings may be indicative of autochthonous transmission of USUV in local passerine populations, although detection of virus in local mosquitoes is required for conclusive confirmation.

Reactive mosquito surveillance conducted in response to this detection confirmed the local presence of *Cx. pipiens*, a confirmed USUV vector in the UK (1), and others known to be competent under experimental conditions (*Cs. annulata)* (13) and to carry USUV in England (14).

The summer of 2025 was the warmest on record in the UK (mean temperature of 16.10°C) (15). Weather conditions were more favourable to mosquitoes (prolonged warm, humid conditions), potentially enhancing conditions for local transmission. However, we cannot yet rule out the possibility that these infected birds acquired USUV elsewhere before migrating to Arran. This year saw the highest recorded number of UK USUV detections to date, with 7.4% (33/446) of bird samples positive compared to 2% (8/397) in 2024. USUV was also detected in birds in Nottinghamshire, Leicestershire, West Yorkshire, Lincolnshire and Lancashire in 2025 (SI File 7), further supporting more general northward expansion in the UK. Sanger or NGS sequencing confirmed USUV lineage Africa 3.2 in all samples.

Results from the enhanced vector surveillance on Arran and wider surveys across the mainland confirm at least 17 mosquito species are present in Scotland, with at least seven considered competent vectors for arboviruses (16). Some UK mosquitoes may be competent for USUV at relatively low temperatures (19C) (13), however local data for Scotland is not yet available and sustained transmission may still be unlikely under typical summer conditions. Further research is needed to assess the likelihood of future USUV outbreaks in Scotland.

## Conclusion

The presence of several known vector species and recent USUV detections in Blackbirds on the Isle of Arran indicate that Scotland is permissive for *Orthoflavivirus* incursion. USUV has been circulating in south-east England for the past 6 years, with no record of human clinical cases so far. The risk to public health from USUV is currently considered very low. Considering climate change and the northward geographic expansion of orthoflaviviruses, there is a clear requirement for improved bird and mosquito surveillance nationwide to better assess risk to animal and human health. Our findings highlight how climate change is increasing the risk of mosquito-borne viruses emerging in higher-latitude countries.

