## Supplementary information for "First Usutu virus detections in wild birds in Scotland, 2025"

**Supplementary material**

***Supplementary Information File 1***- Observations of blackbirds on Isle of Arran, Scotland, between 2023-2025.

| Date | No. of blackbirds | Clinical signs | Location |
|---|---|---|---|
| **04/04/2023** | 1 | unable to fly, head tilt, subsequently euthanised | Not mentioned |
| **April 2024** | 1 | ataxia | Brodick Museum |
| **01/08/2024** | 1 | ataxia, torticollis | Corrie |
| **April 2025** | 1 | ataxia | Brodick Museum |
| **17/06/2025** | 1 | head tilt, unable to fly, subsequently euthanised | Site 1, Brodick |
| **24/06/2025** | 5-6 | ataxic, marked torticollis, wing droop, good general body condition suggesting sudden onset. | Site 1, Brodick |
| **June 2025** | 3 | found dead | Sandbraes Whiting Bay |
| **June 2025** | 2 | found dead | Whiting Bay garden |
| **end of June** | 1 | pronounced feather fluffing | Site 2 Brodick |
| **21/07/2025** | 1 | found dead | Site 3 Brodick |
| **23/07/2025** | 1 | ataxia then subsequently died | Clauchlands Point Lamlash. |
| **24/07/2025** | 3-4 | ataxia then subsequently died | Site 4, Brodick |
| **28/07/2025** | 1 | ataxia then subsequently died | Site 3, Brodick |
| **28/07/2025** | 1 | found dead | Kildonan |
| **29/07/2025** | 1 | Ataxia, feather fluffing, head down. | Site 5, Brodick |

***Supplementary Information File 2-*** Bioinformatic pipeline for next-generation sequencing analysis of the two positive Blackbird samples

Sequencing reads were obtained over a 72-h sequencing window. Adapter reads were removed with Porechop (v0.2.4) and quality checks were performed and sequences trimmed using Chopper (v0.10.0). Host genome removal and reference guided alignment was performed using bowtie2 (v2.5.2) and bcftools (v1.23) and no reads mapped to the USUV consensus sequence used (GenBank accession number: NC006551). *De novo* assembly was performed using MEGAHIT (v1.2.9) and the contigs produced were used to identify viruses through ViralVerify (v1.1) and ViralComplete (v1). The outputs of these programmes then underwent BLAST (v2.5.0) screening using the NCBI “viruses_nt” database (available at: https://ftp.ncbi.nlm.nih.gov/blast/db) to confirm the identity of viral contigs.

***Supplementary Information File 3***- The fifty-nine Usutu virus sequences obtained from GenBank and used in phylogenetic analysis.

| Accession number | Date | Location | Lineage |
|---|---|---|---|
| **MN122238** | 15/08/2018 | Midden Groningen, Netherlands | Africa 3.1 |
| **MN122245** | 22/08/2018 | Westerveld, Netherlands | Africa 3.2 |
| **MN122232** | 07/08/2018 | Tynaarlo, Netherlands | Africa 3.2 |
| **MN122231** | 07/08/2018 | Venlo, Netherlands | Africa 3.2 |
| **MN122246** | 22/08/2018 | Hardenberg, Netherlands | Africa 3.2 |
| **MN122144** | 21/08/2016 | Tytsjerksteradiel, Netherlands | Africa 3.2 |
| **MN122133** | 09/08/2018 | Noordenveld, Netherlands | Africa 3.2 |
| **OR141604** | 2021 | Wildpark Schwarze Berge (NI), Germany | Africa 3.2 |
| **OR141605** | 2021 | Zoo Rostock (MV), Germany | Africa 3.2 |
| **MN122230** | 07/08/2018 | Noordosterpolder, Netherlands | Africa 3.2 |
| **MN122236** | 14/08/2018 | Westland, Netherlands | Africa 3.2 |
| **MN122249** | 23/08/2018 | Lochem, Netherlands | Africa 3.2 |
| **MN122245** | 22/08/2018 | Westerveld, Netherlands | Africa 3.2 |
| **MN122227** | 03/08/2018 | Bilthoven, Netherlands | Africa 3.2 |
| **MN122224** | 02/08/2018 | Ermelo, Netherlands | Africa 3.2 |
| **OR141603** | 2021 | Wildpark Schwarze Berge (NI), Germany | Africa 3.2 |
| **PV779337** | 24/09/2024 | Denmark | Africa 3.2 |
| **PV779328** | 31/08/2024 | Denmark | Africa 3.2 |
| **PV779341** | 10/09/2024 | Denmark | Africa 3.2 |
| **PV779335** | 14/09/2024 | Denmark | Africa 3.2 |
| **MW001216** | 01/07/2020 | London, United Kingdom | Africa 3.2 |
| **OM202464** | 01/09/2021 | London, United Kingdom | Africa 3.2 |
| **MN122248** | 22/08/2018 | Zuidhorn, Netherlands | Africa 3.3 |
| **MN122146** | 06/09/2016 | Andelst, Netherlands | Africa 3.3 |
| **MN122237** | 15/08/2018 | Noordoostpolder, Netherlands | Africa 3.3 |
| **MN122234** | 10/08/2018 | Raalte, Netherlands | Africa 3.3 |
| **MN122247** | 22/08/2018 | Groningen, Netherlands | Africa 3.3 |
| **MN122228** | 06/08/2018 | Middelburg, Netherlands | Africa 3.3 |
| **PV779342** | 11/09/2024 | Denmark | Africa 3.3 |
| **MN122225** | 02/08/2018 | Rotterdam, Netherlands | Africa 3.3 |
| **MN122226** | 03/08/2018 | Leiden, Netherlands | Africa 3.3 |
| **MN122222** | 02/08/2018 | Rijswijk, Netherlands | Africa 3.3 |
| **PV779329** | 13/09/2024 | Denmark | Europe 2 |
| **PV779326** | 10/09/2024 | Denmark | Europe 2 |
| **PV779325** | 01/09/2024 | Denmark | Europe 2 |
| **PV779339** | 21/09/2024 | Denmark | Europe 2 |
| **PV779333** | 17/09/2024 | Denmark | Europe 2 |
| **OR141598** | 2020 | Berlin (BE), Germany | Europe 2 |
| **PV779323** | 30/08/2024 | Denmark | Europe 2 |
| **PV779331** | 14/09/2024 | Denmark | Europe 2 |
| **PV779338** | 15/09/2024 | Denmark | Europe 2 |
| **OR141597** | 2020 | Guben (GB), Germany | Europe 2 |
| **PV779324** | 02/09/2024 | Denmark | Europe 2 |
| **PV779336** | 17/09/2024 | Denmark | Europe 2 |
| **PV779340** | 24/09/2024 | Denmark | Europe 2 |
| **OR141596** | 2020 | Erlau/ Schweikershain (SN), Germany | Europe 2 |
| **PV779334** | 17/09/2024 | Denmark | Europe 2 |
| **PV779332** | 16/09/2024 | Denmark | Europe 2 |
| **OR166137** | 2017 | Mönchengladbach (NW), Germany | Europe 3 |

| | | | |
|---|---|---|---|
| **OR166138** | 2017 | Zoo Dresden (SN), Germany | Europe 3 |
| **OR166136** | 2017 | Soest (NW), Germany | Europe 3 |
| **OR141595** | 2020 | Friedberg (HE), Germany | Europe 3 |
| **OR141594** | 2020 | Wiesbaden (HE), Germany | Europe 3 |
| **OR141593** | 2020 | Limburg (HE), Germany | Europe 3 |
| **PV779327** | 12/09/2024 | Denmark | Europe 3 |
| **MN122229** | 03/08/2018 | Oldambt, Netherlands | Europe 3 |
| **MN122154** | 20/09/2016 | Vlijmen, Netherlands | Europe 3 |
| **MN122152** | 12/09/2016 | Zevenaar, Netherlands | Europe 3 |
| **MN122215** | 11/09/2017 | Eastermar, Netherlands | Europe 3 |

***Supplementary Information File 4*** – Field sampling methods used for mosquitoes and birds during enhanced surveillance on Arran between Oct 2025-Feb 2026:

*i. Mosquito sampling:* Field surveillance for overwintering and host-seeking mosquitoes was conducted on the island between October 2025 and February 2026. Sixteen potential hibernacula, such as outhouses, were initially identified from satellite maps and permission was sought for sampling of overwintering mosquitoes. Permission was also sought for placement of traps at these sites. Additional potential hibernacula were identified opportunistically. Publicly accessible buildings and shelters across the island were thoroughly searched for overwintering mosquitoes. Hand-held battery-powered aspirators were used to collect mosquitoes. Mosquitoes were transferred alive into falcon tubes and placed immediately onto dry ice.

Host-seeking traps (Biogents BG-Pro) baited with carbon dioxide were set up at nine sheltered vegetated locations where access had been granted. Traps were run for 24-48 hours. Collections were placed immediately onto dry ice.

*ii. Bird sampling*:

Field surveillance for overwintering birds was conducted in the same areas of the mosquito sampling, but for birds we relied more heavily on private gardens. We collected a total of 67 blood samples between October 2025 and February 2026, over a total of 10 capture days. Blood samples were taken from a total of 43 different blackbirds. In Brodick, 11 blackbirds in October, 6 blackbirds in December and 12 blackbirds in February were sampled. In Sliddery, 6 blackbirds in January and 4 blackbirds in February were sampled. In Kildonan, 4 blackbirds were sampled in February. We also collected a total of 24 blood samples from an additional 8 passerine bird species spread across the same sites sampled for blackbirds. Blackbirds and other passerines were captured using mist nets and potter traps (under British Trust for Ornithology licences to PJB and DMD). Blood samples were taken from the brachial vein (under Home Office Project Licence PP2193834 to DMD).

***Supplementary Information File 5*** – Laboratory methods used for molecular confirmation of mosquito species, and USUV screening by RT-qPCR (birds and mosquitoes) and serology (birds) from samples collected during enhanced surveillance on Arran between Oct 2025-Feb 2026:

*i. Molecular identification of Cx. pipiens sl samples*: DNA was extracted from individual mosquito legs of morphologically identified cryptic *Culex* mosquitoes using 20 µL squishing buffer, prepared with 10 mM Tris HCl, 1 mM EDTA, 25 mM NaCl and 10 µL proteinase K.

Samples were incubated at 37°C for 20-minutes before proteinase K was inactivated by heating to 95°C for 2-minutes. Extracted DNA was tested using two PCR assays, first to distinguish *Culex pipiens* and *Culex torrentium* using the primers of Smith et al. (2004), then confirming *Culex pipiens* biotype using the primers of Bahnck et al. (2006). Results were confirmed by running products on a 2.5% agarose gel, and bands were visualised using the Biorad GelDoc Go Imaging System.

*ii. RNA extraction, RT and qPCR for Usutu virus screening in mosquitoes and bird blood*: RNA was extracted from individual whole mosquitoes in DNA/RNA shield, bird whole blood in DNA/RNA shield or bird red blood cells placed in DNA/RNA shield after serum removal. Samples were homogenized (Precellys 24, Bertin Technologies) in buffer RLT (Qiagen) at 6500g for 30 sec. RNA was extracted using the RNeasy Mini Kit (Qiagen) following the manufacturer's instructions. In some cases, the volume of reagents was adjusted proportionately according to the volume of bird blood sampled. The recommended DNase I treatment (Qiagen) was also performed. RNA concentration was then measured using a nanodrop and complementary DNAs (cDNAs) were synthesized using the High-Capacity cDNA Reverse Transcription kit (Thermo Fisher Scientific) and using the maximal RNA quantity recommended per reaction following the manufacturer's protocol. qPCR assays were performed using TaqMan Fast Universal PCR Master Mix (2X) (Thermo Fisher Scientific) according to the manufacturer's protocol using previously published USUV primers and probe (1) and using an ABI 7500 Fast PCR machine. A standard curve of a plasmid containing the USUV fragment (Azenta Life Sciences) amplified by PCR was used as positive control. Results were analysed with the 7500 Software v2.0.6.

*iii. Serological testing of bird serum samples for flavivirus exposure:* Serum was recovered from field collected bird blood samples after centrifugation at 11000g for 90s, heat inactivated at 56-60C for 30mins and stored at 4 degrees for no longer than 3 weeks. We used the ID Screen Flavivirus Competition Kit (Innovative Diagnostics, Grabels, France), a commercially available enzyme-linked immunosorbent assay (ELISA), to search for presence of anti-Flavivirus prE antibodies (1). Samples were processed according to manufacturer's protocol.

***Supplementary Information File 6**:* Details of locations where mosquito surveys and bird mist netting was carried out, the number and type of samples collected and the results of Usutu virus testing either by PCR or serology.

**Table Y.** Results of mosquito and bird sampling from sites across the Isle of Arran

| City | Sampling sites | Date | Mosquitoes | Birds/Ring Number | PCR ID | PCR USUV | Serology |
|---|---|---|---|---|---|---|---|
| **Brodick** | Arran Heritage Museum | Oct. | 0 | NA | NA | NA | NA |
| | Forest surrounding Rosaburn Ducks | Oct. | *Aedes detritus* | NA | NA | Negative | NA |
| | | | *Culiseta annulata* | | | Negative | |
| | | Feb. | *Culiseta annulata* | NA | NA | Negative | NA |
| | | | *Culiseta annulata* | NA | NA | Negative | NA |

| | | | | | | |
|---|---|---|---|---|---|---|
| | | *Culiseta annulata* | NA | NA | Negative | NA |
| | | *Culiseta annulata* | NA | NA | Negative | NA |
| Public toilets | Oct. | 0 | NA | NA | NA | NA |
| Site 6, Brodick r | Oct. | 0 | NA | NA | NA | NA |
| Site 2, Brodick | Oct. | 0 | Blackbird LR84156 | NA | Negative | Negative |
| | | | Blackbird LR84157 | | Negative | Negative |
| | | | Blackbird LR84158 | | Negative | Negative |
| | | | Blackbird LR84159 | | Negative | Negative |
| | | | Blackbird LR84160 | | Negative | Negative |
| | | | Blackbird LR84161 | | Negative | Negative |
| | | | Blackbird LR84162 | | Negative | Negative |
| | | | Blackbird LR84163 | | Negative | Negative |
| | | | Blackbird LR84164 | | Negative | Negative |
| | | | Blackbird LR84165 | | Negative | Negative |
| | | | Blackbird LR84166 | | Negative | Negative |
| | | | Song Trush RL79929 | | Negative | Negative |
| Brodick Castle | Jan. | 0 | NA | NA | NA | NA |
| Private property | Dec. | NA | Blackbird LT19250 | NA | Negative | Negative |
| | | | Blackbird LT19251 | | Negative | Negative |
| | | | Blackbird LT19252 | | Negative | Negative |
| | | | Blackbird LT19253 | | Negative | Negative |
| | | | Blackbird LT19254 | | Negative | Negative |
| | | | Blackbird LT19255 | | Negative | Negative |
| | Jan. | 0 | House Sparrow PZ40503 | | Negative | NA |
| | | | House Sparrow PZ40504 | | Negative | Negative |

| | | | | | | | |
|---|---|---|---|---|---|---|---|
| | | | | House Sparrow PZ40505 | | Negative | Negative |
| | | | | House Sparrow PZ40506 | | Negative | Negative |
| | | | | Robin BXH7924 | | Negative | Negative |
| | | | | Robin BXH7926 | | Negative | NA |
| | | | | Chaffinch BXH7925 | | Negative | Negative |
| | | | | Blue Tit BXH7921 | | Negative | Negative |
| | Site 1, Brodick | Feb. | NA | Blackbird LT19260 | NA | Negative | Negative |
| | | | | Blackbird LT19261 | | Negative | Negative |
| | | | | Blackbird LT19262 | | Negative | Negative |
| | | | | Blackbird LT19263 | | Negative | Negative |
| | | | | Blackbird LT19264 | | Negative | Negative |
| | | | | Blackbird LT19265 | | Negative | Negative |
| | | | | Blackbird LT19267 | | Negative | Negative |
| | | | | Blackbird LT19268 | | Negative | NA |
| | | | | Blackbird LT19269 | | Negative | Negative |
| | | | | Blackbird LT19270 | | Negative | Negative |
| | | | | Blackbird LT19271 | | Negative | Negative |
| | | | | Blackbird LT19273 | | Negative | Negative |
| | | | | House Sparrow PZ40529 | | Negative | Negative |
| | | | | House Sparrow PZ40541 | | Negative | Negative |
| | | | | House Sparrow PZ40542 | | Negative | Negative |
| | | | | Robin BXH7960 | | Negative | Negative |
| | | | | Robin BXH7963 | | Negative | NA |
| | | | | Robin BXH7964 | | Negative | Negative |
| | | | | Starling | | Negative | Negative |

| | | | | | | | |
|---|---|---|---|---|---|---|---|
| | | | | LT19266 | | | |
| | | | | Chaffinch BXH7956 | | Negative | Negative |
| **Lamlash** | Middletons Camping | Oct. | *Culex p.* s.l. | NA | *Culex p. pipiens* | Negative | NA |
| **Whiting Bay** | Public Toilets | Oct. | 0 | NA | NA | NA | NA |
| | Private property | Feb. | 0 | NA | NA | NA | NA |
| **Kildonan** | Private property | Feb. | NA | Blackbird LT21332 | NA | Negative | Negative |
| | | | | Blackbird LK67585 | | Negative | Negative |
| | | | | Blackbird LT21333 | | Negative | Negative |
| | | | | Blackbird LK67589 | | Negative | Negative |
| | | | | Great Tit PY16170 | | Negative | Negative |
| | Eas Mor Ecology | Dec. | 0 | NA | NA | NA | NA |
| **Sliddery** | Private property | Jan. | NA | Blackbird LT21311 | NA | Negative | Negative |
| | | | | Blackbird LT21312 | | Negative | Negative |
| | | | | Blackbird LT21313 | | Negative | Negative |
| | | | | Blackbird LT21314 | | Negative | Negative |
| | | | | Blackbird LT21306 | | Negative | Negative |
| | | | | Starling LT21307 | | Negative | Negative |
| | | | | Starling LK67659 | | Negative | Negative |
| | | | | Blackbird LT21315 | | Negative | NA |
| | | | | Dunnock PY46125 | | Negative | Negative |
| | | | | Chaffinch BTJ1032 | | Negative | NA |
| | | Feb. | | Blackbird LT21055 | | Negative | NA |
| | | | | Blackbird LT21082 | | Negative | Negative |
| | | | | Blackbird LT21090 | | Negative | Negative |
| | | | | Blackbird LT21314 | | Negative | Negative |
| | | | | Chaffinch BTJ1064 | | Negative | Negative |

| | | | | | | | |
|---|---|---|---|---|---|---|---|
| **Shiskine** | Bridgend Campsite | Oct. | *Culex p. s.l.* | NA | *Culex p. pipiens* | NA | NA |
| | | | *Anopheles claviger* | | NA | | |
| | Public Toilets | | 0 | | NA | | |
| **Lochranza** | Lochranza Campsite | Oct. | 0 | NA | NA | NA | NA |
| | Public Toilets | | *Culex p. s.l.* | | *Culex p. pipiens* | Negative | |
| | Public Toilets | Feb. | *Culiseta annulata* | NA | NA | Negative | NA |
| | | | *Culiseta annulata* | NA | NA | Negative | NA |
| | | | *Culiseta annulata* | NA | NA | Negative | NA |
| | | | *Culiseta annulata* | NA | NA | Negative | NA |
| | | | *Culiseta annulata* | NA | NA | Negative | NA |
| | | | *Culiseta annulata* | NA | NA | Negative | NA |
| | | | *Culiseta annulata* | NA | NA | Negative | NA |
| **Sannox** | Public Toilets | Oct. | 0 | NA | NA | NA | NA |
| **Glen Rosa** | Camp site Toilets | Oct. | 0 | NA | NA | NA | NA |
| **Corrie** | Caves | Feb. | *Culex p. s.l.* | NA | *Culex p. pipiens* | Negative | NA |
| | | | *Culex p. s.l.* | NA | *Culex p. pipiens* | Negative | |
| | | | *Culex p. s.l.* | NA | *Culex p. pipiens* | Negative | |
| | | | *Culex p. s.l.* | NA | *Culex p. pipiens* | Negative | |
| | | | *Culiseta annulata* | NA | NA | Negative | |
| | | | *Culiseta annulata* | NA | NA | Negative | |
| | | | *Culiseta annulata* | NA | NA | Negative | |
| | | | *Culiseta annulata* | NA | NA | Negative | |
| | | | *Culiseta annulata* | NA | NA | Negative | |
| | | | *Culiseta annulata* | NA | NA | Negative | |

***Supplementary Information File 7-*** Details of the USUV positive samples from Northern England, 2025.

| Location | Grid reference | Bird species | Lineage |
|---|---|---|---|
| Nottinghamshire | SK 6554461423 | Blackbird | Africa 3.2 (confirmed Sanger) |
| Leicestershire | SK 67724081300 | Greenfinch | Africa 3.2 (confirmed NGS) |
| West Yorkshire | SE 1855227264 | Blackbird | Africa 3.2 (confirmed Sanger) |
| Lincolnshire | TF 2017666530 | Blackbird | Africa 3.2 (confirmed Sanger) |
| Lancashire | SD 6293240698 | Common Buzzard | Africa 3.2 (confirmed Sanger) |

***References***


1. Jöst H, Bialonski A, Maus D, Sambri V, Eiden M, Groschup MH, et al. Isolation of usutu virus in Germany. Am J Trop Med Hyg. 2011;85(3):551-3.
